# A Novel Combinatorial Approach Integrating Experimental and Computational Analysis of Antioxidant Activity: Evaluating Catechin and L-Ascorbic Acid in Serum

**DOI:** 10.1101/2024.08.22.609135

**Authors:** Ayeshum Rasool, Chinanu Chidi, Sophie Rigaut, Symone Carty, Chirine Soubra-Ghaoui, Richa Chandra

## Abstract

Cardiovascular disease (CVD) remains the leading cause of morbidity and mortality globally, with oxidative stress playing a pivotal role in its progression. Free radicals produced via oxidative stress contribute to lipid peroxidation, leading to subsequent inflammatory responses, which then result in atherosclerosis. Antioxidants inhibit these harmful effects through their reducing ability, thereby preventing oxidative damage. In this study, we introduce a with computational models simulating hydrophilic and hydrophobic serum environments. We optimized the Ferric Reducing Ability of Plasma (FRAP) assay at a microscale level to evaluate the antioxidant activity of L-ascorbic acid (vitamin C) and catechin, a phytochemical found in green tea, in normal and hypertriglyceridemic serum. Hypertriglyceridemic serum, characterized by increased hydrophobic lipid content, provides a model to examine the impact of serum triglycerides on antioxidant activity. Additionally, we employed computational models using the Gaussian software to simulate the hydrogen atom transfer (HAT) mechanism, calculating free energy changes and bond dissociation energy (BDE) to assess the antioxidant potency of the studied compounds in both hydrophilic and hydrophobic environments. The computational results align with the experimental finding offering a unique combinatorial approach to assess antioxidant activity in both normal and hypertriglyceridemic serum, with potential implications for clinical interventions.

## Introduction

Cardiovascular disease (CVD) is the leading cause of morbidity and mortality in the world.(1) Biologically, oxidative stress through free radicals generated by reactive oxygen species (ROS), reactive nitrogen species, and sulfur is prominent in the development of CVD. At the same time, free radicals are a normal and important part of cellular functions. (2–5) For example, hydrogen peroxide (H_2_O_2_), an ROS that is involved in vascular homeostasis, in excess causes direct lipid peroxidation which can lead to the modification of lipoproteins, which carry lipids in serum, and a subsequent inflammatory response resulting in atherosclerotic lesions.(3) The harmful effects of free radicals are blocked by antioxidants, which are chemicals that prevent oxidation through their ability to reduce ROS for example.(6,7) Common foods such as pomegranates, berries, wine, juices, and bananas all contain antioxidants.(8)

In the work presented here, we introduce a combinatorial approach involving experimental evaluation of antioxidant activity in normal serum and hypertriglyceridemic serum, which has more hydrophobic constituents in the form of lipids, in conjunction with computational models that mimic the hydrophilic and hydrophobic environments of serum. We optimized the Ferric Reducing Ability of Plasma (FRAP) assay (9) at a microscale level, which is useful for sample throughput (10). In contrast to traditional FRAP assays, our experimental method includes antioxidants in the biological environment of serum, which unlike plasma, lacks clotting factors such as fibrinogens as well as EDTA, an antioxidant used in plasma collection procedures. As serum is more deplete in protein content, the effect of hydrophobicity based on triglyceride levels on antioxidant activity can be more closely examined.(11) The FRAP assay works through an iron (III) to iron (II) reduction mechanism, which correlates with oxidation. The reaction is nonspecific, and any half-reaction with a less-positive redox potential will drive the ferric ion reduction.(9). As such, the assay is a useful measure of the antioxidant activity by measuring the electron donating ability of each antioxidant.(12)

L-ascorbic acid or vitamin C, a highly water-soluble antioxidant found in many fruits, and catechin (**Fig 1**), a polyphenolic antioxidant commonly found in green tea and red wine, are evaluated using our two-prong approach.(13–15) Green tea increases antioxidant activity, plays a role in the prevention of cardiovascular disease, and inverses mortality rates.(16,17) In fact, a significant inverse correlation between green tea extract catechin concentration and cholesterol as well as abdominal fat has been previously demonstrated (14,18). L-ascorbic acid is an efficient free radical scavenger which revives the impaired production of endothelium-derived nitric oxide and improves endothelial vasomotor function in arteries lowering the incidence of CVD.(19) Tea flavonols such as catechin are more powerful antioxidants compared to flavonoids such as ascorbic acid and as such are good options to examine the combinatorial approach as a proof of concept.(20,21)

**Fig 1.**
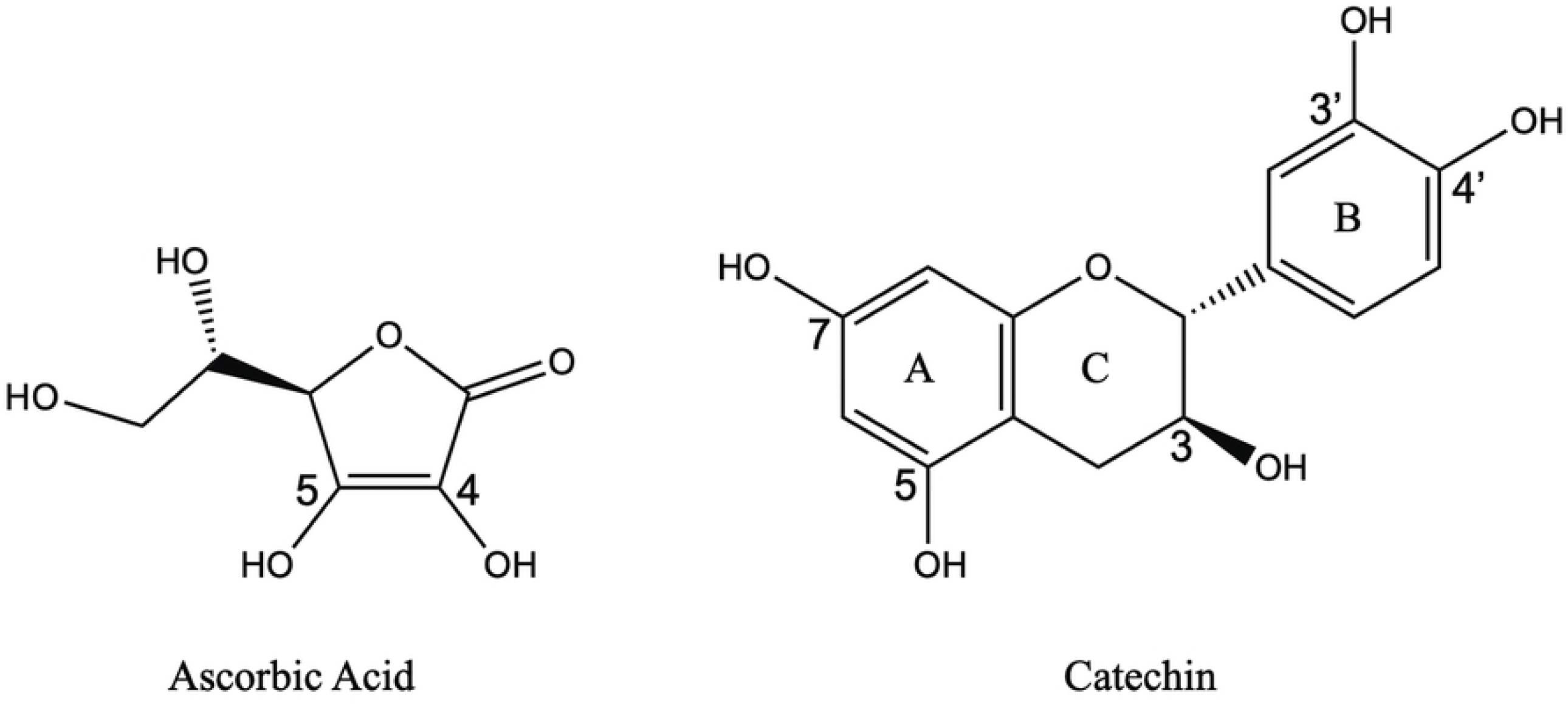
Structures of L-ascorbic acid and catechin with A-, B- and C-rings labeled and with numbered hydroxyl groups involved in the computational hydrogen atom transfer mechanism.

Here we examine the activity of ascorbic acid and catechin experimentally with the FRAP assay in normal triglyceride level serum (<150 mg/dL) and severely hypertriglyceridemic serum (>800 mg/dL). We compare the experimental results with computational models in hydrophilic (water) and hydrophobic (benzene) environments. The severely hypertriglyceridemic samples consist of a higher concentration of nonpolar lipids. We select benzene as our solvent for simplicity and to provide insight into the impact of a pure nonpolar environment on antioxidant activity. The HAT (hydrogen atom transfer) mechanism is one of the most common mechanisms employed for the study of the antioxidant activity of flavonoids.(22) Using the Gaussian (23) software, we calculate free energy changes following the hydrogen atom transfer (HAT) mechanism illustrated in **Equation 1** where AOH represents the antioxidant.

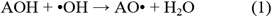

In the HAT reaction, the antioxidant (AOH) loses a hydrogen atom to the highly reactive hydroxide radical resulting in the formation of water and a highly stable antioxidant radical (AO•) that is less reactive. The formation of the most stable radical reflects the greatest theoretical antioxidant activity. Bond dissociation energy (BDE) values are also calculated for the reaction mechanism in Equation 2, which evaluates an antioxidant’s hydrogen donating ability. The lower the BDE value, the greater the H-donating ability and the greater the antioxidant activity.

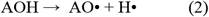

Our combinatorial approach presented here effectively integrates experimental evaluation of antioxidant activity in both normal and hypertriglyceridemic serum—characterized by increased hydrophobic lipid content—with computational models that assess the energetic differences between antioxidants. This combined methodology is highly valuable for studying a wide range of antioxidants in various physiological conditions and their diverse mechanisms. Our novel approach can be leveraged in future studies to offer deeper insights into the physiological activity of antioxidants and potentially guide dietary or clinical interventions.

## Materials and Methods

(+)-Catechin hydrate, L-Ascorbic Acid, Iron (III) Chloride hexahydrate, 2,4,6-Tris(2 pyridyl)-s-triazine, (±)6-Hydroxy-2,5,7,8 tetramethylchroman-2-Carboxylic Acid, sodium acetate•3H_2_0, ethanol, methanol, and hydrochloric acid were all purchased from Sigma Aldrich Co (St. Louis, MO). All clinical serum samples (**Supporting Information: Tables S1 and S2**) with no identifying information were purchased from Discovery Life Sciences, Inc. (Los Osos, CA).

For the experimental assays, aqueous antioxidant solutions were prepared as controls in a 1:4 ratio of water and 40 µM antioxidant in 10% ethanol using a microscale approach of the established protocol of Benzie and Strain (9). For example, 60 µL water and 180 µL of 40 µM catechin or L-ascorbic acid in 10% ethanol were combined. Serum samples were prepared in a similar fashion with a 1:4 ratio of serum (in lieu of water) and 10% ethanol. Briefly, these samples contain 60 µL serum (normal triglyceride values: 57-144 mg/dL, n=11 and severe hypertriglyceridemia: 827-1096 mg/dL, n=13) and 180 µL of 10% ethanol. Serum samples were also combined with antioxidant and prepared at the same ratio of 1:4 for the serum and antioxidant in 10% ethanol. These samples include 60 µL serum and 180 µL of 40 µM catechin or L-ascorbic acid in 10% ethanol. A solution of 180 µL of 10% ethanol and 60 µL water was prepared as the reagent blank. Trolox, a standard vitamin E analog used in FRAP assays, was prepared for calibration at increasing concentrations from 50 µM to 2.5 mM. In short, 60 µL of trolox and 180 µL of 80% methanol were combined as established previously.(9) All the above solutions were incubated at 37 ºC for 1 hour following preparation. A FRAP reaction reagent was prepared with 10 mL of 20 mM FeCl_3_*6H_2_O, 10 mL of 10 mM TPTZ, and 50 mL sodium acetate buffer (pH = 3.6). After the one-hour incubation at 37ºC, the FRAP reagent (1800 µL) was added to all solutions for a 5-minute incubation at 37ºC. We measured the absorbance of all solutions in triplicate at 593 nm at various times following the assay on Biotek’s EPOCH microplate spectrophotometer. The final measurement was taken at 180 minutes when the increase in antioxidant activity became stable for catechin and ascorbic acid. A LINEST calibration (b=0) was performed for each assay, and activities of all controls and samples are described in Trolox equivalents.

In conjunction with experimental assays, computational modeling calculations were performed with Gaussian 16 software. All geometry optimizations were carried out at the m06(24) density functional level of theory employing the triple ζ basis set 6-311++G(d,p)(25)-(26) augmented with diffuse(27) and polarization(28) functions. Vibrational frequencies were computed at the same level of theory to confirm that the optimized geometries are minima and to obtain enthalpy and free energy values. All geometries were also optimized, and frequencies were calculated with solvent effects for water and benzene employing the self-consistent reaction field polarizable conductor model SCRF-CPCM. (29,30)

Stabilization energies are calculated using free energy difference, ΔG, of the products compared to the reactant in hydrogen atom transfer (HAT) reaction represented in **Equation 1**. Bond dissociation energy (BDE) values were calculated as the enthalpy difference at 298 K for the reaction in **Equation 2**. The gas phase served as the control. The OH groups numbered on the antioxidant structures illustrated in **Fig 1** are assessed individually and then in combination using the HAT mechanism for the two calculations.

## Results

The experimental results demonstrate interassay precision based on Trolox standards assayed at the same time as all samples tested with m=(7.3 ± 1.8) × 10^−4^ (LINEST function, b=0, n=12.) All measurements of samples and controls are described in Trolox equivalents (TE). FRAP based measurements of the serum samples without antioxidant provide a level of oxidation that takes place in the serum environment as the ferric ion is reduced and the serum is oxidized (**Table 1)**.

**Table 1.** Oxidation Measurement in Trolox Equivalents of Normal Triglyceride Serum Samples (<150 mg/dL), n=11.

| Sample Number | Triglyceride Concentration (mg/dL) | Serum Oxidation (TE) | SD (TE) |
| --- | --- | --- | --- |
| 1 | 57 | 412 | 3 |
| 2 | 65 | 474 | 11 |
| 3 | 79 | 625 | 2 |
| 4 | 79 | 570 | 9 |
| 5 | 85 | 458 | 4 |
| 6 | 85 | 565 | 6 |
| 7 | 109 | 473 | 3 |
| 8 | 115 | 495 | 3 |
| 9 | 115 | 591 | 3 |
| 10 | 119 | 475 | 2 |
| 11 | 124 | 442 | 2 |
| **Average** | **94** | **507** | **\*17** |

\*All measurements were made in triplicate. The overall standard deviation for the averaged measurements was calculated using the Root Sum of Squares (RSS) method. Specifically, the individual standard deviations of the samples were combined using the equation $RSS = \sum (SD_i^2)$ where $SD_i$ represents the standard deviation of each sample.

When we examine the level of oxidation in the normal triglyceride level samples (<150 mg/dL; average = 94 ± 23 mg/dL), the intrinsic level of oxidation measured by FRAP is 507 ± 17 TE; whereas in the severely hypertriglyceridemic samples (>800 mg/dL; average = 936 ± 84 mg/dL), the intrinsic level of oxidation measured by FRAP trends at a higher average value of 851 ± 23 TE (**Table 2**). These results are expected as increased triglyceride levels are known to be pro-oxidative due to higher concentrations of lipid peroxidation products that can be formed.(31)

**Table 2.** Oxidation Measurement in Trolox Equivalents of Hypertriglyceridemic Triglyceride Serum Samples (>800 mg/dL), n=13.

| Sample Number | Triglyceride Concentration (mg/dL) | Serum Oxidation (TE) | SD (TE) |
| --- | --- | --- | --- |
| 12 | 827 | 868 | 2 |
| 13 | 829 | 1016 | 8 |
| 14 | 868 | 466 | 13 |
| 15 | 868 | 619 | 13 |
| 16 | 868 | 770 | 6 |
| 17 | 921 | 666 | 4 |
| 18 | 927 | 947 | 3 |
| 19 | 965 | 869 | 3 |
| 20 | 975 | 754 | 4 |
| 21 | 983 | 988 | 4 |
| 22 | 994 | 1039 | 3 |
| 23 | 1053 | 1107 | 4 |
| 24 | 1096 | 955 | 2 |
| **Average** | **936** | **851** | **\*23** |

\*All measurements were made in triplicate with SD (standard deviation) reported. The overall standard deviation for the averaged measurements was calculated using the Root Sum of Squares (RSS) method. Specifically, the individual standard deviations of the samples were combined using the equation $RSS = \sum (SD_i^2)$ where $SD_i$ represents the standard deviation of each sample.

When the antioxidants are examined independently, catechin activity averages at 356 ± 27 TE (n=7), whereas ascorbic acid activity averages significantly lower at 10 ± 5 TE (n=7). These results align with previous studies, which reported catechin’s activity to be approximately four times that of Trolox, while ascorbic acid’s activity was close to Trolox’s level.(21)

We lastly examine FRAP activity when the antioxidants were combined with serum. As expected, catechin has a higher activity in normal triglyceride serum compared to ascorbic acid for all samples (**Tables 3**).

**Table 3.** Catechin and Ascorbic Acid FRAP Activities in Trolox Equivalents in Normal Triglyceride Serum Samples (<150 mg/dL), n=11.

| Sample Number | Triglyceride Concentration (mg/dL) | Catechin Activity(TE) | SD (TE) | Ascorbic Acid Activity (TE) | SD (TE) |
| --- | --- | --- | --- | --- | --- |
| 1 | 57 | 424 | 5 | 28 | 6 |
| 2 | 65 | 173 | 12 | 43 | 13 |
| 3 | 79 | 272 | 4 | 57 | 5 |
| 4 | 79 | 252 | 18 | 6 | 21 |
| 5 | 85 | 436 | 6 | 35 | 5 |
| 6 | 85 | 256 | 18 | 8 | 10 |
| 7 | 109 | 255 | 15 | 59 | 5 |
| 8 | 115 | 479 | 8 | 60 | 3 |
| 9 | 115 | 243 | 21 | 8 | 7 |
| 10 | 119 | 600 | 4 | 71 | 3 |
| 11 | 124 | 298 | 9 | 34 | 2 |
| **Average** | **94** | **335** | **\*\*41** | **37** | **\*\*30** |

\*All measurements were made in triplicate with SD (standard deviation) reported. The overall standard deviation for the averaged measurements was calculated using the Root Sum of Squares (RSS) method. Specifically, the individual standard deviations of the samples were combined using the equation $RSS = \sum (SD_i^2)$ where $SD_i$ represents the standard deviation of each sample.

A similar increase in activity is observed in the hypertriglyceridemic serum samples, with catechin showing a more pronounced absolute increase compared to ascorbic acid in all samples. (**Table 4**).

**Table 4.** Catechin and Ascorbic Acid FRAP Activities in Trolox Equivalents in Severely Hypetriglyceride Serum Samples (>800 mg/dL), n=13.

| Sample Number | Triglyceride Concentration (mg/dL) | Catechin Activity(TE) | SD (TE) | Ascorbic Acid Activity (TE) | SD (TE) |
| --- | --- | --- | --- | --- | --- |
| 12 | 827 | 458 | 5 | 84 | 3 |
| 13 | 829 | 736 | 8 | 134 | 8 |
| 14 | 868 | 460 | 15 | 67 | 13 |
| 15 | 868 | 354 | 6 | 36 | 9 |
| 16 | 868 | 226 | 46 | 19 | 12 |
| 17 | 921 | 482 | 5 | 59 | 4 |
| 18 | 927 | 537 | 4 | 153 | 4 |
| 19 | 965 | 552 | 3 | 42 | 5 |
| 20 | 975 | 593 | 10 | 60 | 6 |
| 21 | 983 | 673 | 6 | 157 | 9 |
| 22 | 994 | 576 | 26 | 36 | 11 |
| 23 | 1053 | 857 | 9 | 172 | 5 |
| 24 | 1096 | 710 | 5 | 17 | 3 |
| **Average** | **936** | **555** | **\*\*59** | **80** | **\*\*28** |

\*All measurements were made in triplicate with SD (standard deviation) reported. The overall standard deviation for the averaged measurements was calculated using the Root Sum of Squares (RSS) method. Specifically, the individual standard deviations of the samples were combined using the equation $RSS = \sum (SD_i^2)$ where $SD_i$ represents the standard deviation of each sample.

On average, the absolute increase in catechin’s activity from the normal triglyceride serum to the severely hypertriglyceridemic serum environment goes from 335 ± 41 TE to 555 ± 59 TE. This is an increase of over 200 TE. Ascorbic acid on the other hand, increases from an average of 37 ± 30 TE (normal triglyceride levels <150 mg/dL) to 80 ± 28 TE (severely hypertriglyceridemic levels >800 mg/dL) at an absolute value less than 50 TE.

In the second part of our combinatorial approach, we compared these experimental findings to Gaussian calculations using the radicalized form of each antioxidant (individually assessing each OH group) to find the forms with the lowest free energy (Table 5).

**Table 5:** Relative stabilization free energies (kcal/mol) of Antioxidants and Free Radicals formed from the HAT Reaction Mechanism.

| Antioxidant | Gas Phase | Water | Benzene |
| --- | --- | --- | --- |
| Ascorbic acid (nonradical form) | 0.00 | 0.00 | 0.00 |
| Ascorbic acid 4OH radical | -34.51 | -41.99 | -37.63 |
| Ascorbic acid 5OH radical | -45.99 | -46.71 | -46.28 |
| Ascorbic acid 4OH-5OH radical | -102.84 | -105.15 | -104.51 |
| Catechin (nonradical form) | 0.00 | 0.00 | 0.00 |
| Catechin 3'OH radical | -34.48 | -39.48 | -37.19 |
| Catechin 4'OH radical | -47.21 | -48.51 | -47.66 |
| Catechin 5OH radical | -36.85 | -37.81 | -37.32 |
| Catechin 7OH radical | -34.84 | -37.60 | -35.86 |
| Catechin 3'OH-4'OH-5OH-7OH radical | -138.88 | -155.56 | -145.80 |

\*The most stable forms are highlighted in gray when one OH group is radicalized.

For ascorbic acid, when the OH group in position 4 is radicalized (**Fig 1**) the corresponding radical, ascorbic acid-4OH is formed. Likewise, when the OH groups in positions 4 and 5 are both radicalized (**Fig 1**) the corresponding radical, ascorbic acid 4OH-5OH is formed (**Table 5**). By radicalizing only one hydroxyl group, the 5OH radical is the most stable form in gas, water and benzene. In the gas phase the 5OH radical is more stable by 11.48 kcal/mol than its 4OH counterpart. By radicalizing both hydroxyl groups at positions 4 and 5, we obtain a greater stabilization energy of −102.84 kcal/mol.

For catechin, the most stable singly radicalized form is catechin-4’OH. When considering the most stable radical form of each antioxidant in the gas phase, we find the catechin 4’OH radical is more stable at −47.21 kcal/mol compared to the ascorbic acid-5OH radical at −36.85 kcal/mol. We observe the same pattern in water and benzene. By radicalizing all four hydroxyl groups at positions 3’, 4’, 5, and 7 we obtain a much greater stabilization energy of −138.88 kcal/mol. The increased stabilization obtained when multiple hydroxyl groups are radicalized indicates that the scavenging activity of the antioxidant is improved as the number of hydroxyl groups attached to the aromatic ring increases. This is in line with the experimental findings that also illustrate an enhanced activity for catechin in all samples compare to L-ascorbic acid. When assessing the stability of the same antioxidant radical in the gas phase compared to its stabilization in water and benzene solvents, we find the greatest stability to occur in water.

According to calculated bond dissociation energy (BDE) values **(Table 6)**, where a lower value reflects greater H-donating ability, we find the H abstraction from the 4’ position on catechin is the most facile with BDE values of 70.48 kcal/mol in the gas phase, 70.28 kcal/mol in water and 70.57 kcal/mol in benzene. The ascorbic acid-5OH radical comes in second with BDE values of 71.22 kcal/mol, 72.99 kcal/mol, and 71.99 kcal/mol in the gas phase, water, and benzene, respectively. In agreement with radical stability assessed through free energy (**Table 5**), we find that the 5OH radical of ascorbic acid reflects the greatest H-donating ability in gas, water and benzene (**Table 6**). Likewise, the catechin 4’-OH radical reflects the greatest H-donating ability (**Table 6**) and forms the most stable radical (**Table 5**).

**Table 6:**
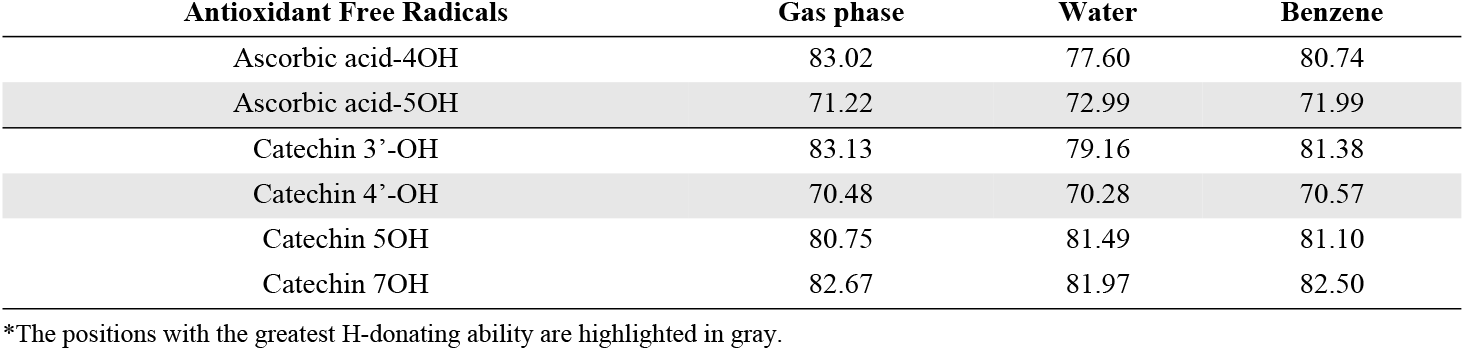
Bond dissociation energies (kcal/mol) for ascorbic acid and catechin free radicals in the gas phase, water, and benzene.

| Antioxidant Free Radicals | Gas phase | Water | Benzene |
| --- | --- | --- | --- |
| Ascorbic acid-4OH | 83.02 | 77.60 | 80.74 |
| Ascorbic acid-5OH | 71.22 | 72.99 | 71.99 |
| Catechin 3'-OH | 83.13 | 79.16 | 81.38 |
| Catechin 4'-OH | 70.48 | 70.28 | 70.57 |
| Catechin 5OH | 80.75 | 81.49 | 81.10 |
| Catechin 7OH | 82.67 | 81.97 | 82.50 |

\*The positions with the greatest H-donating ability are highlighted in gray.

## Discussion

Catechin’s activity is enhanced in severely hypertriglyceridemic serum samples as evidenced through the FRAP assay data. Though catechin’s structure includes several hydroxyl groups which generally increase hydrophilicity, the presence of the 7OH group on the A ring (Fig 1) assists in the formation of a six-membered aromatic ring that in turn increases the hydrophobicity of the molecule.(32) It would be interesting to explore the effect of hypertriglyceridemia on other antioxidants (i.e. quercetin and ficetin) that have a similar structure and hydrophobicity using the FRAP assay and the computational models. Serum oxidation increases in severely hypertriglyceridemic samples and illustrates the potential for using the microscale FRAP assay as a measurement of oxidative susceptibility, as there is a need to develop the total oxidant status that are not time-consuming or costly.(33)

The computational and experimental data collectively demonstrate that catechin exhibits significantly greater antioxidant activity than ascorbic acid. This superiority is attributed to catechin’s structural configuration, which includes a higher number of hydroxyl groups that benefit from resonance stabilization via the hydrogen atom transfer (HAT) mechanism. The antioxidant efficacy of catechins is intricately linked to the position and quantity of these hydroxyl groups, with a reduction in antioxidant capacity corresponding to fewer hydroxyl groups on catechin’s B ring. (34) The computational analysis (Table 5) further reveals that both ascorbic acid and catechin radicals show enhanced stability in aqueous environments, though the hydrogen-donating ability remains relatively unaffected by different phases—gas, benzene, or water. While benzene alone does not adequately mimic the serum environment where antioxidants are likely to partition between the aqueous blood medium and lipid components (35), the computational findings offer valuable insights into the structural and energetic advantages of catechin over ascorbic acid.

The FRAP assay proves to be a facile, robust, and reproducible tool, offering critical insights into the comparative antioxidant activities of various compounds. Its application is particularly valuable in assessing the influence of serum triglycerides on antioxidant efficacy, especially for polyphenolic structures that can partition between aqueous and lipid environments. Given that triglycerides contribute to oxidative mechanisms within the physiological milieu of blood, other mechanisms can be experimentally tested and compared with the computational data. Additionally, investigating other antioxidants with varied structural characteristics, such as quercetin or epicatechin, could provide a broader understanding of antioxidant behavior under different physiological conditions such as hypercholesterolemia. Moreover, exploring the interactions with other blood constituents like copper (36) and nitric oxide (37), or considering variations in serum composition due to factors like diet, disease, or age, could yield deeper insights into how these variables influence oxidative mechanisms and antioxidant efficacy. Collectively, the integration of the FRAP assay with computational models opens new avenues for exploring the nuanced mechanisms behind antioxidant activity, offering potential pathways for clinical and dietary interventions.

## Supporting information

Additional information about normolipidemic serum samples including triglyceride results (mg/dL) is available in **Tables S1** and **S2** along with the computation model’s Cartesian coordinates of the HAT complexes in the gas phase in **Table S3**.

